# Forty-four years of change: Capercaillie and Black grouse responses to changes in forestry, climate and vole decline in central Sweden

**DOI:** 10.64898/2026.04.28.721321

**Authors:** Tomas Willebrand, Rolf Brittas, Jonas Kindberg

## Abstract

Human land use has greatly affected the natural landscapes in most parts of the world, and Scandinavian industrial forestry has substantially altered the age structure of the boreal forest over the last 75 years. In this study, we analyzed 44 years of line transect counts of Capercaillie and Black grouse collected in an industrial forest landscape in central Sweden between 1980 and 2024. The area was heavily logged in the 1960s and 1970s, followed by large-scale replanting. A hierarchical Gompertz state–space model to assess long-term population dynamics of both species. The adult state model included a brood production submodel, and forest age structure, snow depth, vole abundance, and spring frost were covariates. Our results reveal strong density dependence and temporally variable environmental effects. Capercaillie and Black grouse exhibited contrasting decadal trends: Capercaillie increased markedly during the early study years, whereas Black grouse declined steeply before recovering later in the series. Increasing proportions of forest aged 21–40 years had a clearly positive effect on Capercaillie, whereas snow depth in the previous winter negatively influenced Capercaillie but not Black grouse. Brood production exhibited substantial interannual unexplained variation, although late spring frost reduced brood size in Capercaillie. Toward the end of the study, the two species reached similar levels of latent adult abundance, demonstrating that both can persist at sustainable densities across a broader range of forest structures than suggested in earlier studies.

## Introduction

Human land use and exploration have changed the natural landscapes in many parts of the world. Industrial forestry is economically important in Scandinavia and has significantly modified the boreal conifer landscape (Esseen et al. 1997). However, forest management has undergone significant and sometimes rapid changes (Jacobsson et al. 2025), and the implementation of revised regeneration practices take decades to come into effect. In the period from 1950s to 1980s, forestry focused on investment in harvest and regeneration. The increased use of clear-cutting practiced reduced the proportion of older forests and created landscapes dominated by young, even-aged stands, potentially affecting specific successional stages-dependent species.

Three forest-dwelling grouse species occur in Scandinavia: Capercaillie *Tetrao urogallus*, Black grouse *Lyrurus tetrix*, and hazel grouse *Tetrastes bonasia*. Early studies of Capercaillie and Black grouse in industrial forests formed the basis of the forestâĂŞsuccession hypothesis: Capercaillie were considered dependent on mature conifer forests, whereas Black grouse were associated with younger stands, heather moorland, or tundra edges (Seiskari, 1962; Swenson and Angelstam, 1993). These studies were conducted during the period when the forest consisted largely of either old stands or recent clear-cuts and plantations (Rolstad and Wegge, 1987; Lande et al. 2014; Finne et al. 2000; Lindén et al. 2000; Angelstam, 2004).

The last decades have seen increased efforts to restore structures that are important for biodiversity and an increase in old and large trees. (Jacobsson et al. 2025). Clear-cut sizes are now restricted, and the proportions of seedling, sapling, pole, and mature stands have become more similar due to long-term planning and rotation (*The Swedish National Forest Inventory* 2025). Recent research has questioned the strong preference of Capercaillie for mature forests (Rolstad, Rolstad, et al. 2007; Sirkiä et al. 2010; Wegge and Rolstad, 2011).

Grouse have historically been a valuable game species in Scandinavia, although harvesting today is primarily recreational. In the period 2019 - 2021, about twenty thousand Capercaillie were harvested annually in Sweden. Harvest statistics show large annual, often cyclic, fluctuations driven largely by nest loss and chick survival (Jahren et al. 2016; Kurki et al. 1997; Wegge, Moss, et al. 2022). Predation, influenced by vole abundance as an alternative prey, is considered a key proximate driver of annual brood variation (Marcstrom, Kenward, et al. 1988; Wegge and Storaas, 1990; Linden, 1988; Small et al. 1993). Other factors, including the onset of spring, weather, and early insect availability, also affect brood success (Tornberg et al. 2012; Willebrand, 1992; Wegge and Kastdalen, 2008; Helminen and Viramo, 1962; Ruottinen et al. 2024; Storaas et al. 2000).

Long-term data are essential for detecting weak but persistent population trends (Lindenmayer et al. 2012; Franklin, 1989). Harvest statistics have often been used to track long-term trends (Afton and Anderson, 2001; Aebischer, 2019; Hjeljord and Loe, 2022), but their reliability for grouse has been questioned (Ranta et al. 2008; Willebrand, Hörnell-Willebrand, et al. 2011).

In this study, we analyzed a unique 44-year long-term time series of line transect counts of Capercaillie and Black grouse. Using a hierarchical stateâĂŞspace model allowed us to separate latent abundance from observation noise, improving inference on population trends, species comparisons, and environmental drivers. Adult abundance and brood production were modeled using covariates related to vole abundance, snow depth, spring frost, and forest age structure. This study aimed to describe key population characteristics and long-term changes in Capercaillie and Black grouse inhabiting an industrial forest landscape.

## Methods

### Study area

The study was conducted within the 32 km^2^ Boda Wildlife Research Station from 1980 to 2024. The landscape consists of commercially managed forest at the border between the southern and middle boreal zones, 20 km inland from the Gulf of Bothnia (61°N, 16°E). See figure 1. Dominant tree species are Scots pine *Pinus sylvestris*, Norway spruce *Picea abies*, and Birch *Betula* spp. The ground vegetation is dominated by Bilberry *Vaccinium myrtillus*, Heather *Calluna vulgaris*, and Wavy-hair grass *Deschampsia flexuosa*. Predators include Goshawk *Accipiter gentilis*, Ural owl *Strix uralensis*, Hooded crow *Corvus corone*, Raven *Corvus corax*, Eurasian jay *Garrulus glandarius*, Red fox *Vulpes vulpes*, Pine marten *Martes martes*, and Stoat *Mustela erminea*.

**Figure 1.**
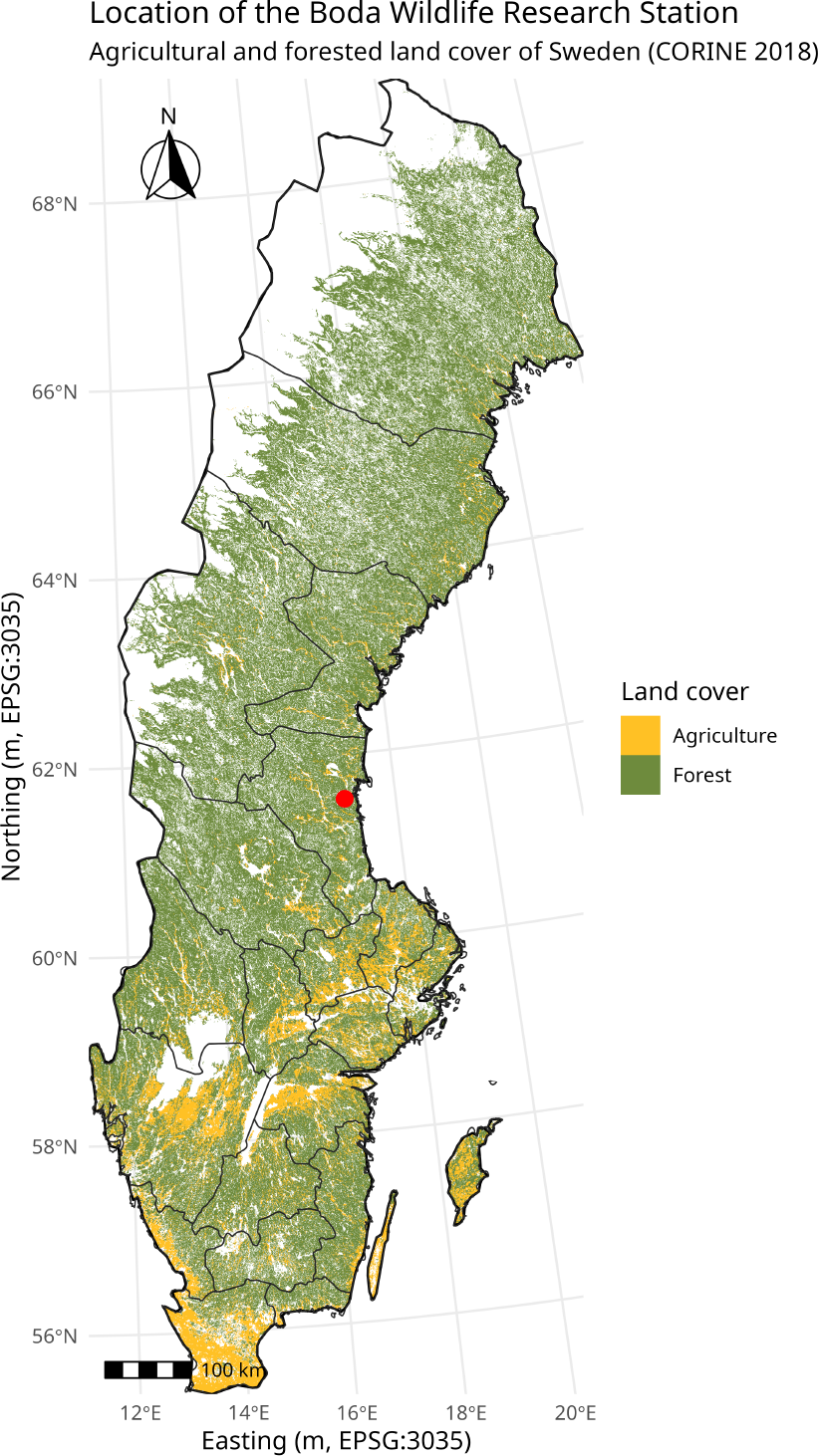
Map of Sweden showing the location of the study area (red dot).

**Figure 2.**
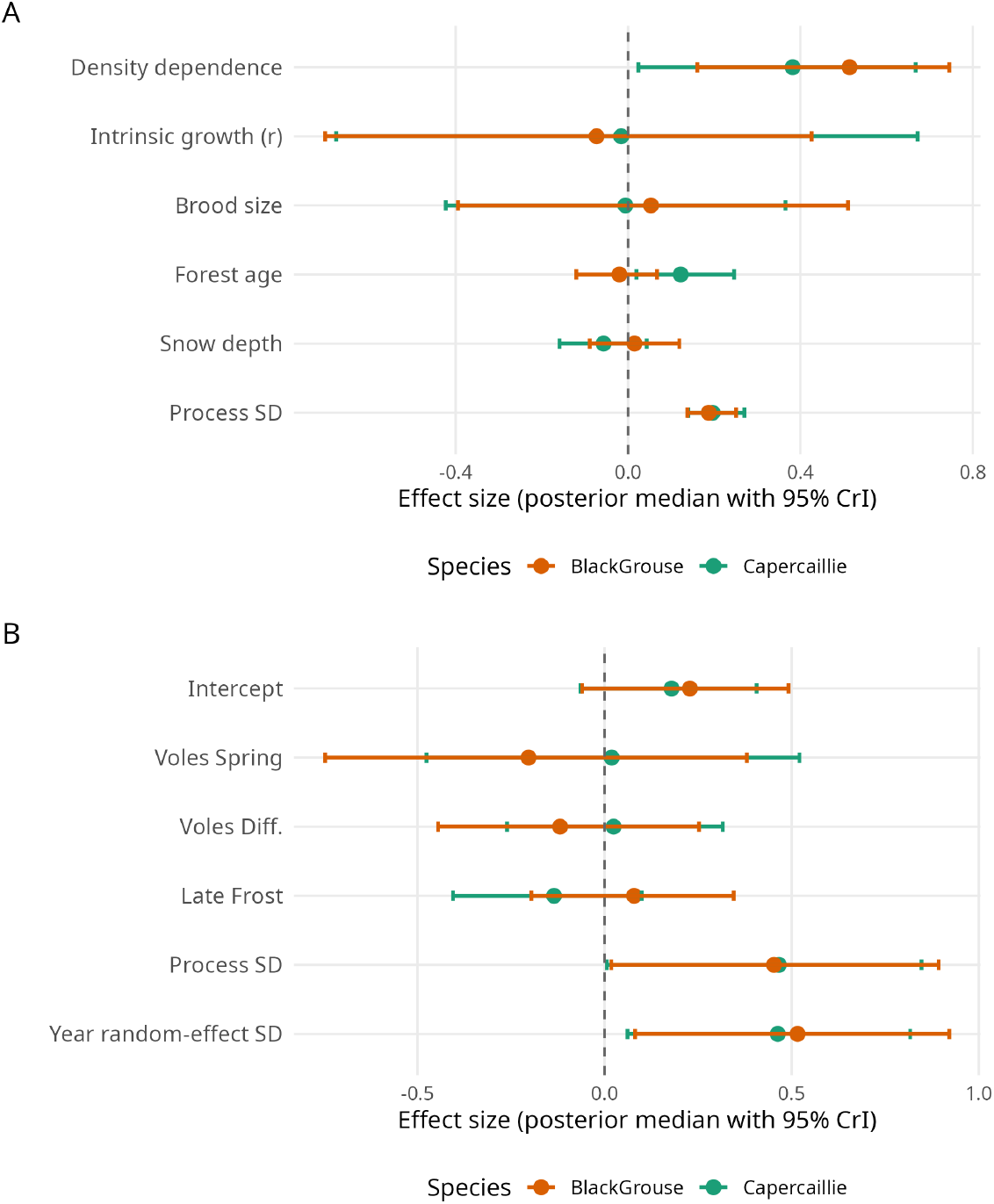
Posterior summaries for the adult and brood model parameters.

**Figure 3.**
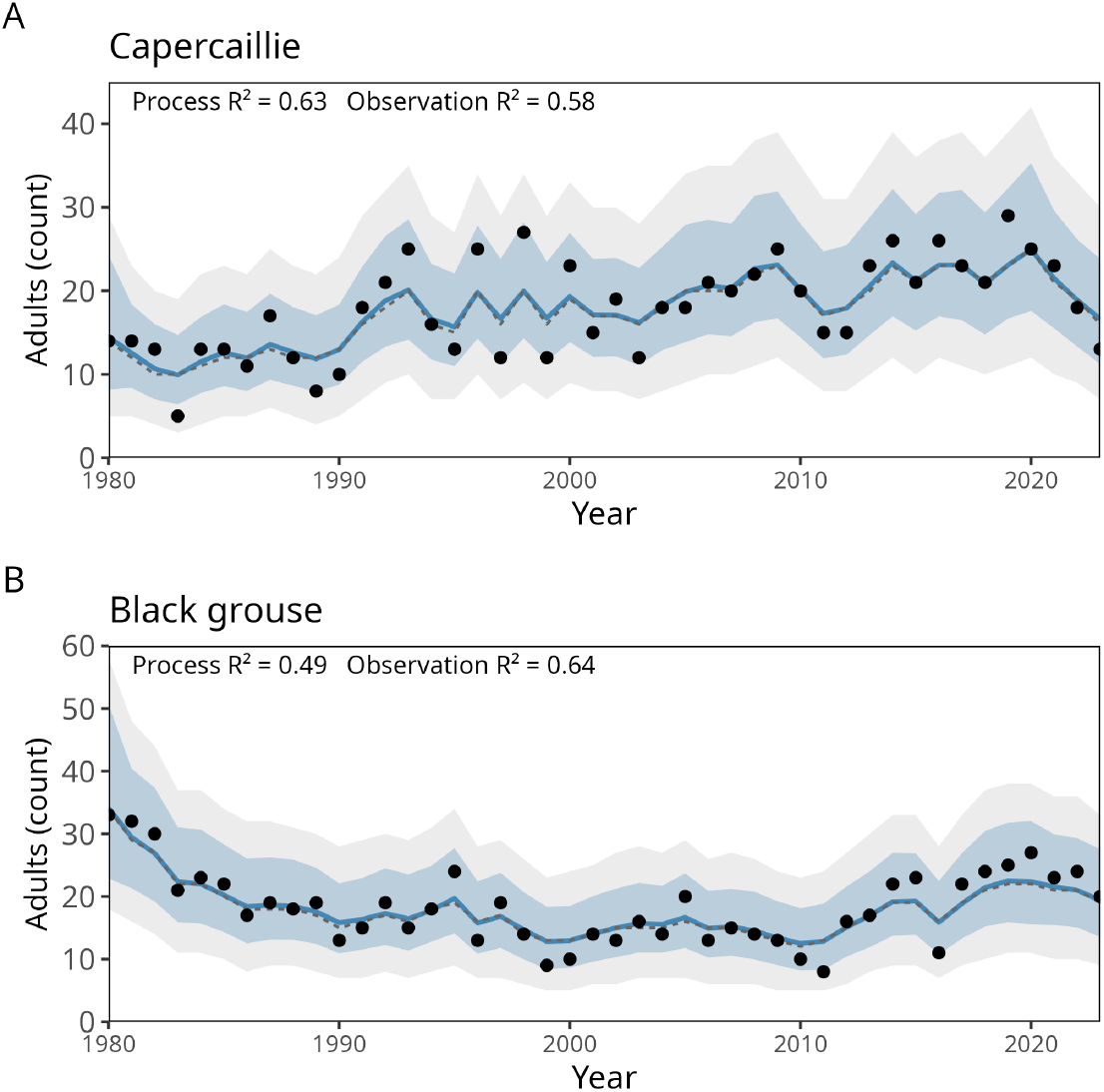
Latent adult abundance (*x*_*t*_) with 95% credible intervals and observed counts.

### Data collection

In 1980, permanent transect lines spaced 400 m apart were established in 1980, with a total length of 84.4 km. Three observers walked 20 m apart, each recording birds within 10 m of their line, yielding a strip width of 60 m (Brittas and Karlbom, 1990; Helle and Lindström, 1991) and a surveyed area of 5.06 km^2^. Surveys were conducted in July when chicks were old enough to be detected. A pilot study suggested that the detection probabilities of adults not accompanied were 54–58% for Capercaillie and 61–64% for Black grouse (Brittas and Karlbom, 1990). Hazel grouse detection was unreliable and was excluded. However, without a more detailed study on the factors affecting detectability, we choose not to model the uncertainty of detection, and therefore, the estimates of latent abundance are an underestimate.

Vole abundance was measured using four 100 m trap lines in May and August (Marcstrom, Hoglund, et al. 1990). Spring numbers used in brood models declined after the early 1980s, and formerly pronounced vole cycles collapsed. The time series of spring numbers can be found in the supplementary materials. Spring numbers rather than autumn numbers were selected because it was expected to affect the breeding success the most, but we also included the difference between spring and autumn catches as a covariate.

Climate data were obtained from the Järvsö meteorological station 42 km west of Boda. Covariates included mean January–February snow depth and the last date in May with temperature below *−*2.2°C (Schwartz and Chen, 2002).

Forest age data were unavailable for the study site; therefore, we used regional statistics from 16,276 km^2^ surrounding the area. The logit-transformed proportion of forest aged 21–40 years captured the major structural shift during the study period (*The Swedish National Forest Inventory* 2025).

All covariates were standardised to zero mean and unit variance. The environmental covariates showed no substantial collinearity, and only date of the last frost was correlated with year.

### State–space model structure

We used hierarchical Bayesian state–space models to separate demographic processes from observation error. To avoid inflating the process variance due to outliers, we used a Student-*t* formulation for the process error. The model consisted of: (1) A Gompertz process model for adult abundance log *x*_*t*_, and (2) A submodel for brood production and chick observations. We report Pr(*β* > 0 | data) for covariate effects.

### Adult abundance model

Capercaillie dynamics were well described by near-Gaussian deviations, whereas Black grouse dynamics exhibited occasional abrupt increases or declines, justifying a heavier-tailed process for that species.

Adult abundance *x*_*t*_ followed a Gompertz process,

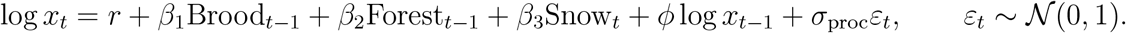

To allow heavy tails, we used a scale-mixture representation with *w*_*t−*1_ ∼ Gamma(*ν*_proc_/2, *ν*_proc_/2), yielding a Student-*t* process with *ν*_proc_ degrees of freedom.

### Observation model for adult counts

The observed adult counts followed a negative binomial distribution to account for overdispersion:

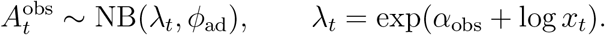

Posterior predictive checks used Freeman–Tukey discrepancies and log-likelihood contributions.

### Brood productivity model

Brood size *B*_*t*_ (chicks per observed female) followed

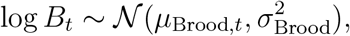

with

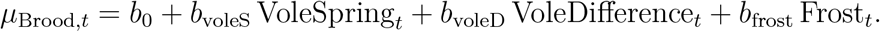

The expected chick counts were *λ*_Young,*t*_ = *B*_*t*_*F*_*t*_.

### Observation model for chicks

The capercaillie chick counts followed a negative binomial distribution. Black grouse chick counts required a zero-inflated negative binomial model because of structural zeros from brood failure.

### Prior distributions

To allow occasional extreme annual changes without inflating overall variability, process noise in the adult state equation was modelled using a Student–*t* innovation implemented via a gamma–normal mixture. The scale parameter *σ*_proc_ received a weakly informative log-normal prior with a lower truncation at 0.08 to prevent unrealistically small process variance and ensure biologically plausible year-to-year fluctuations. This specification stabilised the estimation, reduced confounding with observation error, and preserved the separation between the typical variation and rare extreme years. The regression coefficients had 𝒩 (0, 1) or 𝒩 (0, 4) priors. Density dependence used *φ* ∼ Beta(3, 3). Process noise and overdispersion parameters used log-normal priors. Black grouse zero inflation used *ψ* ∼ Beta(2, 2).

### Model validation

Models were fitted in JAGS using three chains, a burn-in of 7,000 iterations, and 18,000 posterior iterations with thinning of two. Convergence was evaluated using traceplots, 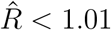, and effective sample sizes. Posterior predictive checks, process and observation *R*^2^, and residual plots were used to assess model adequacy. As recommended in (Gelman and Shalizi, 2013), we interpret these values directly without additional confidence intervals.

### Rolling decadal trends

We estimated decade-scale trends by fitting regressions of log *x*_*t*_ on standardised time within rolling 10-year windows across the posterior draws. The annual percentage change was computed as exp(*β*_trend_) *−* 1. This approach provids a time–resolved assessment of long–term population phases.

### Forest structure and species ratio

To visualise species responses to forest structure, we examined

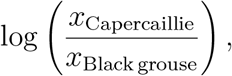

and related this ratio to the proportion of forest aged 21–40 years using a Gaussian GAM fitted to posterior mean ratios. The smooth is based on posterior means; uncertainty around the relationship is therefore somewhat larger than the confidence band alone suggests.

## Results

### Adult population model

The posterior estimates indicated strong density dependence and species-specific environmental responses (Table 2). For Capercaillie, forest age had a clearly positive effect (posterior probability 0.99), and snow depth had a predominantly negative effect (0.871). The effects of variation in brood size were weak (0.518). For Black grouse, the posterior probabilities for brood size, forest age, and snow depth were all below 0.70. The process variance was moderate for both species, and the Student-*t* process allowed for occasional larger deviations.

**Table 1.** Definitions of symbols and parameters used in the state–space model.

| Symbol / Parameter | Definition |
| --- | --- |
| $x_t$ | Latent adult abundance at time $t$ |
| $A_t^{\text{obs}}$ | Observed adult count in year $t$ |
| $B_t$ | Latent brood size (chicks per female) |
| $Y_t$ | Observed chick count in year $t$ |
| $F_t$ | Observed number of adult females |
| $r$ | Intrinsic growth rate (Gompertz process) |
| $\beta_{1\text{Brood}_{t-1}}$ | Effect of mean brood size in year $t - 1$ |
| $\beta_{2\text{Forest}_{t-1}}$ | Effect of forest proportion aged 21–40 years |
| $\beta_{3\text{Snow}_t}$ | Effect of snow depth in year $t$ |
| $\phi$ | Strength of density dependence ($0 < \phi < 1$) |
| $\sigma_{\text{proc}}$ | Process standard deviation (adult model) |
| $\nu_{\text{proc}}$ | Degrees of freedom of the Student-$t$ process |
| $\alpha_{\text{obs}}$ | Adult observation intercept |
| $\phi_{\text{ad}}$ | NB dispersion parameter (adult counts) |
| $b_0$ | Brood model intercept |
| $b_{\text{voleS}}$ | Effect of spring vole index |
| $b_{\text{voleD}}$ | Effect of spring–autumn vole difference |
| $b_{\text{frost}}$ | Effect of late spring frost |
| $\sigma_{\text{Brood}}$ | Process SD for brood size |
| $\sigma_y$ | Year random-effect SD (brood model) |
| $\phi_{\text{Young}}$ | NB dispersion for chick counts |
| $\psi$ | Zero-inflation probability (Black grouse only) |

**Table 2.** Posterior summaries for parameters in the adult Gompertz state model. Values are presented as posterior medians with 95% credible intervals.

| Definition | Parameter | Capercaillie |  |  | Black grouse |  |  |
| --- | --- | --- | --- | --- | --- | --- | --- |
|  |  | Median | 2.5% | 97.5% | Median | 2.5% | 97.5% |
| Intrinsic growth | $r$ | -0.016 | -0.677 | 0.672 | -0.073 | -0.703 | 0.426 |
| Brood size | $\beta_1$ | -0.006 | -0.423 | 0.365 | 0.053 | -0.395 | 0.510 |
| Forest age | $\beta_2$ | 0.122 | 0.019 | 0.246 | -0.020 | -0.120 | 0.067 |
| Snow depth | $\beta_3$ | -0.057 | -0.159 | 0.043 | 0.015 | -0.089 | 0.119 |
| Density dependence | $\phi$ | 0.382 | 0.024 | 0.667 | 0.514 | 0.161 | 0.745 |
| Process SD | $\sigma_{\text{proc}}$ | 0.196 | 0.140 | 0.270 | 0.187 | 0.138 | 0.251 |
| Dispersion (NB) | $\phi_{\text{ad}}$ | 60.041 | 29.660 | 121.537 | 65.668 | 28.856 | 157.061 |

### Brood model

The effects of vole abundance and frost are summarised in Table 3. Only late spring frost showed a clear negative effect on Capercaillie brood size (posterior probability 0.876). Brood sizes exhibited substantial year-to-year variation, captured by process and annual random-effect parameters. The chick overdispersion parameters indicated moderate extra-Poisson variability.

**Table 3.** Posterior summaries for the brood model parameters. Values are presented as posterior medians with 95% credible intervals.

| Definition | Parameter | Capercaillie |  |  | Black grouse |  |  |
| --- | --- | --- | --- | --- | --- | --- | --- |
|  |  | Median | 2.5% | 97.5% | Median | 2.5% | 97.5% |
| Intercept | $\beta_0$ | 0.179 | -0.064 | 0.406 | 0.228 | -0.060 | 0.492 |
| Voles Spring | $\beta_1$ | 0.019 | -0.476 | 0.521 | -0.203 | -0.747 | 0.380 |
| Voles Diff. | $\beta_2$ | 0.024 | -0.261 | 0.316 | -0.119 | -0.445 | 0.252 |
| Late Frost | $\beta_3$ | -0.135 | -0.405 | 0.099 | 0.079 | -0.196 | 0.345 |
| Process SD | $\sigma_{\text{Brood}}$ | 0.466 | 0.006 | 0.847 | 0.453 | 0.018 | 0.893 |
| Year random-effect SD | $\sigma_y$ | 0.463 | 0.061 | 0.817 | 0.516 | 0.082 | 0.921 |
| Dispersion (NB) | $\phi_{\text{Young}}$ | 55.731 | 10.085 | 276.767 | 25.105 | 3.097 | 197.349 |

### Latent abundance dynamics

The latent abundance trajectories showed substantial temporal variation for both species. Posterior credible intervals reflected process uncertainty beyond the sampling noise. Species showed markedly different population phases, consistent with the decadal trend analyses summarised below.

### Decadal trends in abundance

Rolling 10-year trends revealed contrasting long-term dynamics (Table 4). Capercaillie increased strongly early in the study and remained largely stable thereafter, with indications of a recent decline. Black grouse declined early but showed considerable recovery in later decades. Variation among rolling windows demonstrates the importance of multi-year context when interpreting long-term trends. Figure 4 shows the time–resolved trend trajectories for both species.

**Table 4.** Summary of rolling decadal trends for the full study period. Values are presented as posterior medians with 95% credible intervals.

| Species | Min trend (%) | Max trend (%) |
| --- | --- | --- |
| Capercaillie | -6.90 (-18.57, 5.53) | 18.19 (2.43, 37.18) |
| Black grouse | -19.25 (-30.08, -7.18) | 19.50 (3.38, 39.79) |

**Figure 4.**
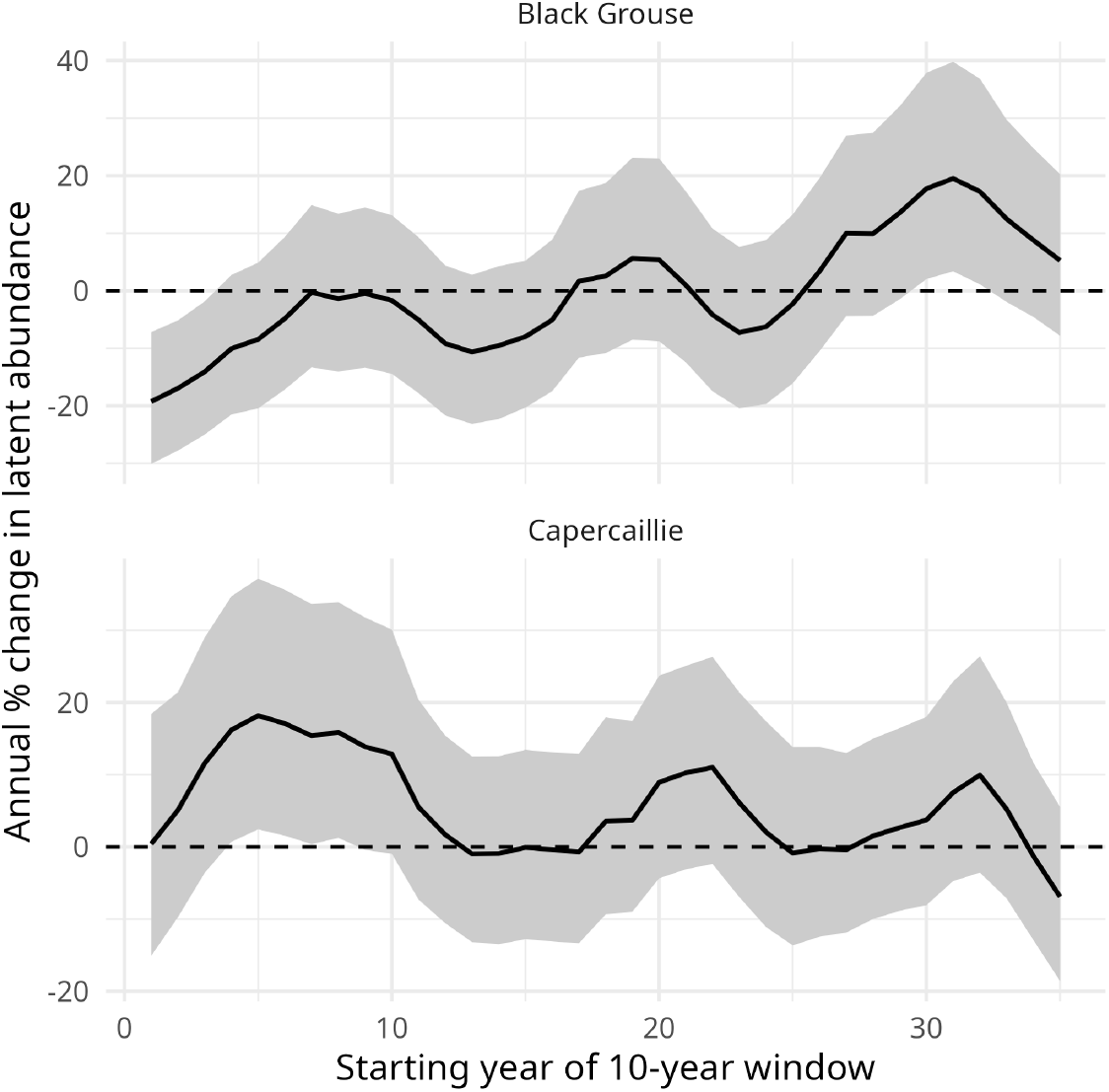
Rolling 10-year trends for Capercaillie and Black grouse.

**Figure 5.**
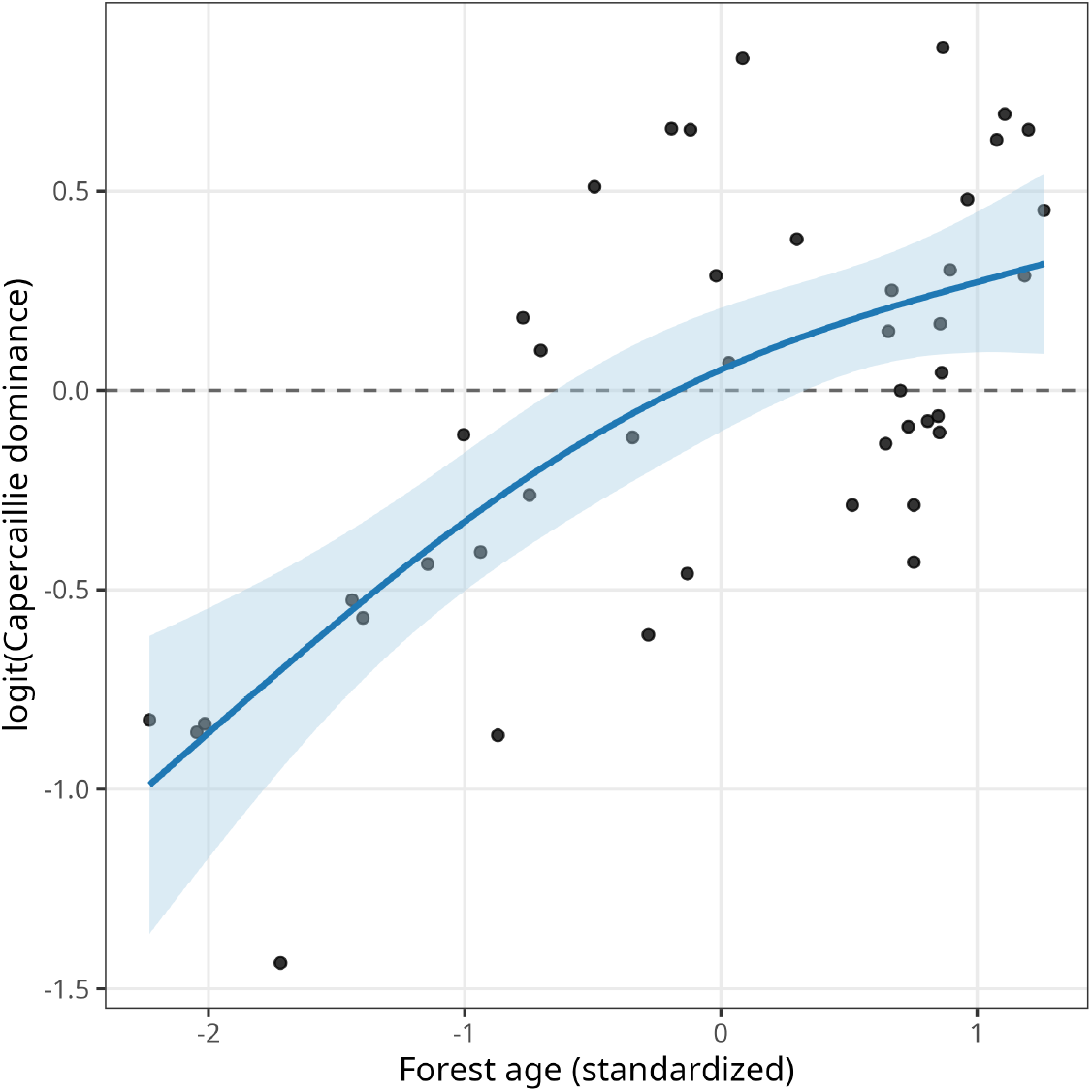
Relationship between forest age structure and log abundance ratio, log(*x*_cap_*/x*_black_).Points are the posterior means and the line shows a GAM smooth with 95% confidence bands.

### Forest age structure and relative dominance

The log abundance ratio exhibited a strong nonlinear relationship with the forest structure. Black grouse dominated when young forest was prevalent, whereas Capercaillie became dominant as the proportion of 21–40-year-old stands increased.

## Discussion

By separating ecological process variation from observation noise, our state–space analysis provided a clearer understanding of long-term population dynamics. In the early years, Capercaillie increased strongly, whereas Black grouse declined. These contrasting patterns coincided with the maturation of forest stands following large-scale clear-cutting in the preceding decades. Toward the end of the study, both species reached similar latent abundances, demonstrating that Capercaillie and Black grouse can persist at sustainable levels across a wider range of forest structures than previously assumed.

Sarcoptic mange entered the Swedish Red fox population for the first time between 1975-1977, and drastically reduced the population within three to five years (Willebrand, Samelius, et al. 2022). This had large effects on several prey populations of the Red fox Lindström et al. 1994, but the Red fox population rapidly recovered from 1985 onwards. In addition to the change in forest structure, we suggest that the initial decline of the Black grouse is also dependent on increased predation after the return of the Red fox, which had been almost absent from the food web for a number of years. We expect a stronger response in Black grouse compared to Capercallie because Black grouse has a faster life-history with lower survival and larger clutches compared to the larger Capercaillie (Jahren et al. 2016).

The strong negative density dependence in both species likely reflects the functional responses of predators within a complex boreal predator–prey system (Lindström et al. 1994; Wallgren et al. 2009; Linden, 1988; Small et al. 1993; Korpimärki and Krebs, 1996). However, density dependence in annual counts should be interpreted as aggregate demographic compensation, not only as evidence for predator effects; it may also integrate variation in survival, dispersal, and recruitment over spatial scales larger than the study area. Forest age structure integrates several ecological mechanisms, including habitat composition, predator distributions, and plant community changes Widenfalk and Weslien, 2009, and thus provides a useful phenomenological index of grouse habitat quality rather than a single causal driver.

Although brood production varied greatly from year to year, its direct effect on adult population change was weaker than expected. Low chick sample sizes, especially for Black grouse, and recruitment processes operating over spatial scales larger than the study area likely reduced the link between local brood production and subsequent adult abundance. Nevertheless, the brood submodel improved identifiability by constraining plausible recruitment variation within the state–space framework.

The alternative prey hypothesis predicts improved grouse reproduction during vole peaks (Hagen, 1952; Angelstam et al. 1984; Breisjøberget et al. 2018; Marcstrom, Kenward, et al. 1988), but the disappearance of vole cycles (Ims et al. 2008) appears to have weakened this mechanism in recent decades (Wegge, Moss, et al. 2022). Late spring frost reduced brood size in Capercaillie, consistent with the importance of early spring vegetation quality for income breeders (Brittas, 1988; Stephens et al. 2009). A very late spring in 1985 postponed Black grouse breeding by more than 10 days (Willebrand, 1988), illustrating the sensitivity of grouse reproduction to spring phenology. Deep midwinter snow may also indirectly influence the conditions. The Capercaillie was negatively affected by snow depth, and contrary to the Black grouse, the snow was not deep enough for the Capercaillie to hide in snow burrows.

Several limitations should be acknowledged. Annual counts cannot resolve within-season demographic processes or short-lived events that influence survival or reproduction. The environmental covariates were broad-scale indices, and finer-scale forestry and weather information might explain the additional variation. The Student-*t* process captures the presence of extreme years but does not attribute them to specific mechanisms. Long-term datasets may also include slow changes in detectability and survey conditions. Despite these limitations, this study provides important insights into grouse ecology and highlights the importance of separating brood production from adult abundance when analyzing population dynamics.

## Conclusions

Rolling decadal trends provided a clear overview of long-term population phases and underscored the importance of continuous monitoring. Using state–space models to separate process and observation variance yielded new insights into the contrasting longterm dynamics linked to forest structure. Future work incorporating details on spatial-scale dispersal and recruitment and finer-scale forest metrics will be essential for understanding the mechanisms driving grouse population changes in the bor4al forest.

## Supporting information

Summary of models and details on results.

## Acknowledgements

We are grateful to the late Dr. N. Höglund, who foresaw the importance of long-term counts and initiated this monitoring programme. The Swedish Hunters’ Association has provided continued support and data curation of the counts since they were initiated.

## Supplementary Materials

Supplementary material can be found in *supplement*.*pdf*.

## References

Aebischer, N. J. (2019). “Fifty-Year Trends in UK Hunting Bags of Birds and Mammals, and Calibrated Estimation of National Bag Size, Using GWCTs National Gamebag Census”. European Journal of Wildlife Research 65.4, p. 64. doi: 10.1007/s10344-019-1299-x.

Afton, A. D. and M. G. Anderson (2001). “Declining Scaup Populations: A Retrospective Analysis of Long-Term Population and Harvest Survey Data”. The Journal of Wildlife Management 65.4, pp. 781–796. doi: 10.2307/3803028. JSTOR: 3803028.

Angelstam, P., E. Lindström, and P. Widén (1984). “Role of Predation in Short-Term Population Fluctuations of Some Birds and Mammals in Fennoscandia”. Oecologia 62.2, pp. 199–208.

Angelstam, P. (2004). “Habitat Thresholds and Effects of Forest Landscape Change on the Distribution and Abundance of Black Grouse and Capercaillie”. Ecological Bulletins 51, pp. 173–187. JSTOR: 20113307.

Breisjøberget, J. I., M. Odden, P. Wegge, B. Zimmermann, and H. Andreassen (2018). “The Alternative Prey Hypothesis Revisited: Still Valid for Willow Ptarmigan Population Dynamics”. PLOS ONE 13.6, e0197289. doi: 10.1371/journal.pone.0197289.

Brittas, R. (1988). “Nutrition and Reproduction of the Willow Grouse Lagopus Lagopus in Central Sweden”. Ornis Scandinavica 19.1, p. 49. doi: 10.2307/3676527. JSTOR: 3676527.

Brittas, R. and M. Karlbom (1990). “A Field Evaluation of the Finnish 3-Man Chain: A Method for Estimating Forest Grouse Numbers and Habitat Use”. Ornis Fennica 67.1 (1), pp. 18–23.

Esseen, P.-A., B. Ehnström, L. Ericson, and K. Sjöberg (1997). “Boreal Forests”. Ecological Bulletins 46, pp. 16–47. JSTOR: 20113207.

Finne, M. H., P. Wegge, S. Eliassen, and M. Odden (2000). “Daytime Roosting and Habitat Preference of Capercaillie Tetrao Urogallus Males in Spring - the Importance of Forest Structure in Relation to Anti-Predator Behaviour”. Wildlife Biology 6.4, pp. 241–249. doi: 10.2981/wlb.2000.022.

Franklin, J. F. (1989). “Importance and Justification of Long-Term Studies in Ecology”. Long-Term Studies in Ecology. Ed. by G.E. Likens. New York, NY: Springer New York, pp. 3–19. doi: 10.1007/978-1-4615-7358-6_1.

Gelman, A. and C. R. Shalizi (2013). “Philosophy and the practice of Bayesian statistics”. British Journal of Mathematical and Statistical Psychology 66.1, pp. 8–38.

Hagen, Y. (1952). “Rovfuglene Og Viltpleien [Raptors and Game Management]”. Oslo: Gyldendal Norsk Forlag.

Helle, P. and J. Lindström (1991). “Censusing Tetranoids by the Finnish Wildlife Triangle Method: Principles and Some Applications”. Ornis Fennica 68.4 (4), pp. 148–157.

Helminen, M. and J. Viramo (1962). “Animal Food of Capercaillie (Tetrao Urogallus) and Black Grouse (Lyrurus Tetrix) in Autumn”. Ornis Fennica 39.1, pp. 1–12.

Hjeljord, O. and L. E. Loe (2022). “The Roles of Climate and Alternative Prey in Explaining 142 Years of Declining Willow Ptarmigan Hunting Yield”. Wildlife Biology 2022.6, e01058. doi: 10.1002/wlb3.01058.

Ims, R., J. Henden, and S. Killengreen (2008). “Collapsing Population Cycles”. Trends in Ecology & Evolution 23.2, pp. 79–86. doi: 10.1016/j.tree.2007.10.010.

Jacobsson, J., J. Fridman, A.-L. Axelsson, and P. Milberg (2025). “An Aging Population? A Century of Change among Swedish Forest Trees”. Forest Ecology and Management 580, p. 122509. doi: 10.1016/j.foreco.2025.122509.

Jahren, T., T. Storaas, T. Willebrand, P. Fossland Moa, and B.-R. Hagen (2016). “Declining Reproductive Output in Capercaillie and Black Grouse 16 Countries and 80 Years”. Animal Biology 66.3–4, pp. 363–400. doi: 10.1163/15707563-00002514.

Korpimärki, E. and C. J. Krebs (1996). “Predation and Population Cycles of Small Mammals”. BioScience 46.10, pp. 754–764. doi: 10.2307/1312851.

Kurki, S., P. Helle, H. Lindén, and A. Nikula (1997). “Breeding Success of Black Grouse and Capercaillie in Relation to Mammalian Predator Densities on Two Spatial Scales”. Oikos 79.2, pp. 301–310. doi: 10.2307/3546014. JSTOR: 3546014.

Lande, U. S., I. Herfindal, T. Willebrand, P. F. Moa, and T. Storaas (2014). “Landscape Characteristics Explain Large-Scale Variation in Demographic Traits in Forest Grouse”. Landscape Ecology 29.1, pp. 127–139. doi: 10.1007/s10980-013-9960-3.

Linden, H. (1988). “Latitudinal Gradients in Predator-Prey Interactions, Cyclicity and Synchronism in Voles and Small Game Populations in Finland”. Oikos (Copenhagen, Denmark), pp. 341–349.

Lindén, H., P. I. Danilov, A. N. Gromtsev, P. Helle, E. V. Ivanter, and J. Kurhinen (2000). “Large-Scale Forest Corridors to Connect the Taiga Fauna to Fennoscandia”. Wildlife Biology 6.3, pp. 179–188. doi: 10.2981/wlb.2000.007.

Lindenmayer, D. B., G. E. Likens, A. Andersen, D. Bowman, C. M. Bull, E. Burns, C. R. Dickman, A. A. Hoffmann, D. A. Keith, M. J. Liddell, A. J. Lowe, D. J. Metcalfe, S. R. Phinn, J. Russell-Smith, N. Thurgate, and G. M. Wardle (2012). “Value of Long-Term Ecological Studies”. Austral Ecology 37.7, pp. 745–757. doi: 10.1111/j.1442-9993.2011.02351.x.

Lindström, E. R., H. Andrén, P. Angelstam, G. Cederlund, B. Hörnfeldt, L. Jäderberg, P.-A. Lemnell, B. Martinsson, K. Sköld, and J. E. Swenson (1994). “Disease Reveals the Predator: Sarcoptic Mange, Red Fox Predation, and Prey Populations”. Ecology 75.4, pp. 1042–1049. doi: 10.2307/1939428.

Marcstrom, V., R. E. Kenward, and E. Engren (1988). “The Impact of Predation on Boreal Tetraonids during Vole Cycles: An Experimental Study”. The Journal of animal ecology, pp. 859–872.

Marcstrom, V., N. Hoglund, and C. J. Krebs (1990). “Periodic Fluctuations in Small Mammals at Boda, Sweden from 1961 to 1988”. The Journal of Animal Ecology 59.2, p. 753. doi: 10.2307/4893. JSTOR: 4893.

Ranta, E., J. Lindström, H. Lindén, and P. Helle (2008). “How Reliable Are Harvesting Data for Analyses of Spatio-Temporal Population Dynamics?” Oikos 117.10, pp. 1461– 1468. doi: 10.1111/j.0030-1299.2008.16879.x.

Rolstad, J., E. Rolstad, and P. Wegge (2007). “Capercaillie Tetrao Urogallus Lek Formation in Young Forest”. Wildlife Biology 13.1, pp. 59–67. doi: 10.2981/0909-6396(2007)13[59:CTULFI]2.0.CO;2.

Rolstad, J. and P. Wegge (1987). “Distribution and Size of Capercaillie Leks in Relation to Old Forest Fragmentation”. Oecologia 72.3, pp. 389–394. doi: 10.1007/BF00377569.

Ruottinen, K. M., M. Melin, J. Miettinen, M. Kervinen, V.-M. Pakanen, J. T. Forsman, and S. Rytkönen (2024). “Assessing the Effects of Drainage and Forest Structure on Presence and Absence of Fledglings of Boreal Grouse”. Global Ecology and Conservation 54, e03150. doi: 10.1016/j.gecco.2024.e03150.

Schwartz, M. and X. Chen (2002). “Examining the Onset of Spring in China”. Climate Research 21, pp. 157–164. doi: 10.3354/cr021157.

Seiskari, P. (1962). “On the Winter Ecology of the Capercaillie, Tetrao Urogallus, and the Black Grouse, Lyrurus Tetrix, in Finland”. Pap Game Res 22, pp. 1–119.

Sirkiä, S., A. Lindén, P. Helle, A. Nikula, J. Knape, and H. Lindén (2010). “Are the Declining Trends in Forest Grouse Populations Due to Changes in the Forest Age Structure? A Case Study of Capercaillie in Finland”. Biological Conservation 143.6, pp. 1540–1548. doi: 10.1016/j.biocon.2010.03.038.

Small, R. J., V. Marcström, and T. Willebrand (1993). “Synchronous and Nonsynchronous Population Fluctuations of Some Predators and Their Prey in Central Sweden”. Ecography 16.4, pp. 360–364. JSTOR: 3683120.

Stephens, P. A., I. L. Boyd, J. M. McNamara, and A. I. Houston (2009). “Capital Breeding and Income Breeding: Their Meaning, Measurement, and Worth”. Ecology 90.8, pp. 2057– 2067. doi: 10.1890/08-1369.1.

Storaas, T., P. Wegge, and L. Kastdalen (2000). “Weight-Related Renesting in Capercaillie Tetrao Urogallus”. Wildlife Biology 6.4, pp. 299–303. doi: 10.2981/wlb.2000.030.

Swenson, J. E. and P. Angelstam (1993). “Habitat Separation by Sympatric Forest Grouse in Fennoscandia in Relation to Boreal Forest Succession”. Canadian Journal of Zoology 71.7, pp. 1303–1310. doi: 10.1139/z93-180.

The Swedish National Forest Inventory (2025). SLU.SE. url: https://www.slu.se/en/Collaborative-Centres-and-Projects/the-swedish-national-forest-inventory/about-the-nfi/ (visited on 01/14/2025).

Tornberg, R., V. Reif, and E. Korpimäki (2012). “What Explains Forest Grouse Mortality: Predation Impacts of Raptors, Vole Abundance, or Weather Conditions?” International Journal of Ecology 2012.1, p. 375260. doi: 10.1155/2012/375260.

Wallgren, M., R. Bergström, K. Danell, and C. Skarpe (2009). “Wildlife Community Patterns in Relation to Landscape Structure and Environmental Gradients in a Swedish Boreal Ecosystem”. Wildlife Biology 15.3, pp. 310–318. doi: 10.2981/08-045.

Wegge, P., R. Moss, and J. Rolstad (2022). “Annual Variation in Breeding Success in Boreal Forest Grouse: Four Decades of Monitoring Reveals Bottom-up Drivers to Be More Important than Predation”. Ecology and Evolution 12.10, e9327. doi:10.1002/ece3.9327.

Wegge, P. and J. Rolstad (2011). “Clearcutting Forestry and Eurasian Boreal Forest Grouse: Long-term Monitoring of Sympatric Capercaillie Tetrao Urogallus and Black Grouse T. Tetrix Reveals Unexpected Effects on Their Population Performances”. Forest Ecology and Management 261.9, pp. 1520–1529. doi: 10.1016/j.foreco.2011.01.041.

Wegge, P. and T. Storaas (1990). “Nest Loss in Capercaillie and Black Grouse in Relation to the Small Rodent Cycle in Southeast Norway”. Oecologia 82.4, pp. 527–530. doi: 10.1007/BF00319796.

Wegge, P. and L. Kastdalen (2008). “Habitat and Diet of Young Grouse Broods: Resource Partitioning between Capercaillie (Tetrao Urogallus) and Black Grouse (Tetrao Tetrix) in Boreal Forests”. Journal of Ornithology 149.2, pp. 237–244. doi: 10.1007/s10336-007-0265-7.

Widenfalk, O. and J. Weslien (2009). “Plant Species Richness in Managed Boreal ForestsEffects of Stand Succession and Thinning”. Forest Ecology and Management 257.5, pp. 1386–1394. doi: 10.1016/j.foreco.2008.12.010.

Willebrand, T. (1992). “Breeding and Age in Female Black Grouse Tetrao Tetrix”. Ornis Scandinavica 23.1, p. 29. doi: 10.2307/3676424. JSTOR: 3676424.

Willebrand, T., M. Hörnell-Willebrand, and L. Asmyhr (2011). “Willow Grouse Bag Size Is More Sensitive to Variation in Hunter Effort than to Variation in Willow Grouse Density”. Oikos 120.11, pp. 1667–1673. doi: 10.1111/j.1600-0706.2011.19204.x.

Willebrand, T., G. Samelius, Z. Walton, M. Odden, and J. Englund (2022). “Declining Survival Rates of Red Foxes Vulpes Vulpes during the First Outbreak of Sarcoptic Mange in Sweden”. Wildlife Biology, wlb3.01014. doi: 10.1002/wlb3.01014.

Willebrand, T. R. (1988). “Demography and Ecology of a Black Grouse (Tetrao Tetrix L.) Population.” PhD thesis, Uppsala University, Sweden.

