## Supplementary material for "Forty-four years of change: Capercaillie and Black grouse responses to changes in forestry, climate and vole decline in central Sweden": Summary of models and details on results.

### Model summary

##### State Variable

- $x_t$: latent adult abundance at year $t$ (Capercaillie or Black grouse).
- $B_t$: latent brood size (chicks per female) in year $t$.

##### Process Model

**Adult abundance (Gompertz state equation):**

$$\log x_t = r + \beta_1 \text{Brood}_{t-1} + \beta_2 \text{Forest}_{t-1} + \beta_3 \text{Snow}_t + \phi \log x_{t-1} + \eta_t,$$

where the innovation term $\eta_t$ follows a Student–$t$ distribution, implemented via a gamma–normal scale mixture.

$$\eta_t = \frac{\sigma_{\text{proc}}}{\sqrt{w_t}} z_t, \qquad z_t \sim \mathcal{N}(0,1), \quad w_t \sim \text{Gamma}(\nu_{\text{proc}}/2, \nu_{\text{proc}}/2).$$

**Brood productivity:**

$$\log B_t \sim \mathcal{N}(\mu_{\text{Brood},t},\, \sigma^2_{\text{Brood}}),$$

$$\mu_{\text{Brood},t} = b_0 + b_{\text{voleS}}\, \text{VoleSpring}_t + b_{\text{voleD}}\, \text{VoleDifference}_t + b_{\text{frost}}\, \text{Frost}_t.$$

##### Observation Model

**Adult counts:** Negative binomial

$$A_t^{\text{obs}} \sim \text{NB}(\lambda_t,\, \phi_{\text{ad}}), \qquad \lambda_t = \exp(\alpha_{\text{obs}} + \log x_t).$$

**Chicks:**

$$Y_t \sim \begin{cases} 0, & \text{with probability } \psi, \\ \text{NB}(\lambda_t^{\text{Young}},\, \phi_{\text{Young}}), & \text{with probability } 1-\psi, \end{cases} \qquad \lambda_t^{\text{Young}} = B_t F_t.$$

##### Priors

- Regression coefficients: $\mathcal{N}(0,1)$ or $\mathcal{N}(0,4)$.
- Density dependence: $\phi \sim \text{Beta}(3,3)$.
- Process SD: $\log \sigma_{\text{proc}} \sim \mathcal{N}(\log \sigma_{\text{mean}}, \log \sigma^2_{\text{sd}})\, T(\log 0.08, \infty)$.
- NB dispersions: log-normal.
- Zero-inflation (black grouse): $\psi \sim \text{Beta}(2,2)$.

### R-scripts and jags code

The R-script forthe Capercaillie and Black grouse state-spaces models can be found in *Run_Gompertz_models.r*. Note that priors are passed to the models as data.

The jags-model can be found in *Gompertz_Full_Option3_NB.jags* for capercaillie and *Gompertz_Full_Option3_ZINB.jags* for black grouse. A directed acyclic graph of the model is shown in figure 1.

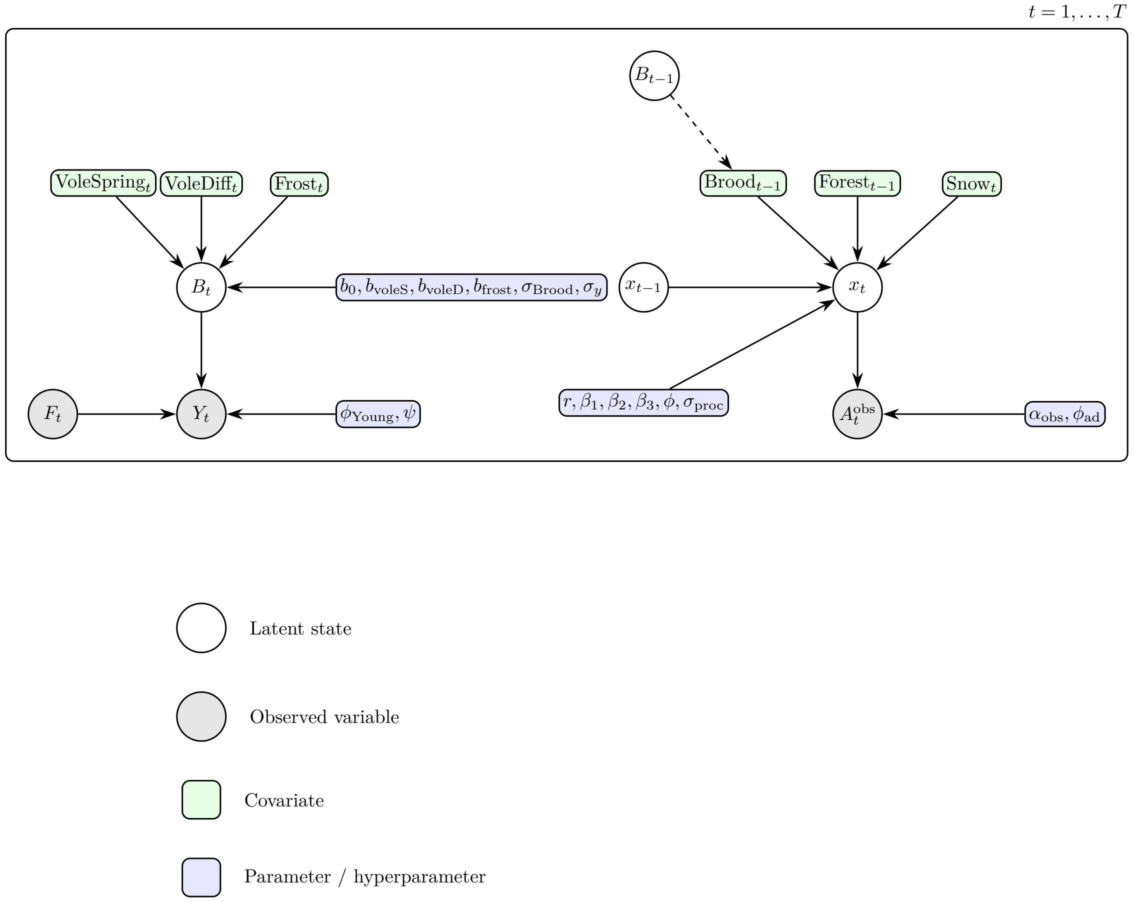


Figure 1: A directed acyclic graph (DAG) visualization of a state-space model used to analyze the demographics of capercaillie and black grouse.

**Prior for process variation.** Annual population changes in the adult state equation were allowed to follow a Student–$t$ distribution to accommodate occasional large deviations from the expected Gompertz dynamics. We implemented the $t$ distribution through a gamma–normal scale mixture, in which the process innovation is written as

$$\eta_t = \frac{\sigma_{\text{proc}}}{\sqrt{w_t}} z_t, \qquad z_t \sim \mathcal{N}(0,1),\ w_t \sim \text{Gamma}(\nu_{\text{proc}}/2,\ \nu_{\text{proc}}/2),$$

so that smaller values of the degrees of freedom parameter $\nu_{\text{proc}}$ generate heavier tails. This parameterisation separates the "typical" year-to-year fluctuation, governed by the scale parameter $\sigma_{\text{proc}}$, from rare and extreme deviations associated with the heavy-tailed mixing distribution.

To ensure biologically plausible levels of annual variation and to prevent the process variance from collapsing towards zero—a common identifiability issue in state–space models—we assigned a weakly informative prior on the logarithmic scale,

$$\log \sigma_{\text{proc}} \sim \mathcal{N}(\log \sigma_{\text{mean}},\, \log \sigma_{\text{sd}}^2)\; T(\log 0.08,\, \infty)\,,$$

where the lower truncation corresponds to a minimum biologically reasonable annual standard deviation. This log-normal formulation restricts $\sigma_{\text{proc}}$ to positive values, provides appropriate right-skewed support, and avoids the need for arbitrary upper bounds. Together with the gamma–mixture representation of the Student–$t$ innovations, this specification allowed the model to capture both moderate annual fluctuations and occasional extreme years without inflating the overall level of process noise.

#### Validation

We initially checked R–hat and effective sample size to confirm model convergence. All R-hat values were below 1.05 and sample sizes were larger than 100. Bayesian p- vales are shown in Table 1

Table 1: Bayesian p-values, Freeman - Tukey PPC for adults & young.

| Species | Age | Bayesian p-value |
|---|---|---|
| Capercaillie | Adult | 0.782 |
| | Young | 0.546 |
| Black grouse | Adult | 0.881 |
| | Young | 0.571 |

The residuals are pictured in figure 2.For adults, most residuals fall between $-0.5$ and $+0.5$, with no systematic bias. The NB dispersion was sufficient and no funnel was evident. We expect that detection variability can explain the few large residuals ($\pm 1$ SD). For young, residuals mostly fall between $-0.5$ and $+0.5$, which indicates a good match between predicted [brood size $\times$ number of females] and actual chick counts. The slight upward skew at larger fitted values is probably explained by a few unusually large broods. There was no obvious curvature (nonlinearity).

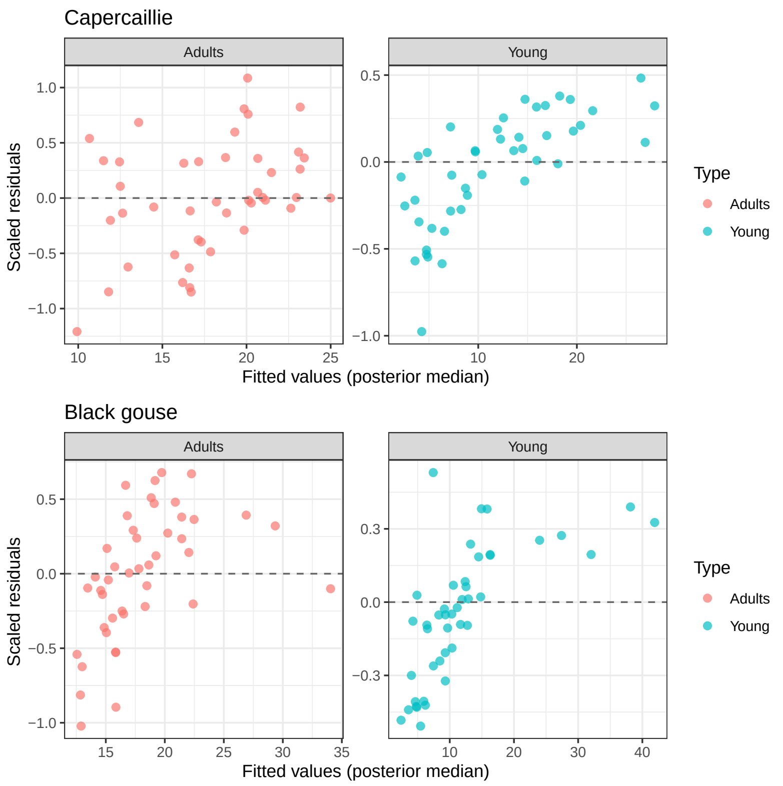


Figure 2: Residual vs fitted plots for Capercaillie and Black grouse.

### Detailed table of decadal trends

Table 2: Decadal trends (annual percent change) for each 10–year rolling window. Posterior medians with 95% credible intervals.

| Start year | End year | Median (%) | 95% CrI |
|---|---|---|---|
| | | Capercaillie | |
| 1 | 10 | 0.420 | -15.073 – 18.412 |
| 2 | 11 | 5.104 | -9.723 – 21.430 |
| 3 | 12 | 11.583 | -3.592 – 28.998 |
| 4 | 13 | 16.205 | 0.708 – 34.732 |
| 5 | 14 | 18.186 | 2.431 – 37.180 |
| 6 | 15 | 17.087 | 1.558 – 35.628 |
| 7 | 16 | 15.392 | 0.377 – 33.651 |
| 8 | 17 | 15.878 | 1.246 – 33.905 |
| 9 | 18 | 13.871 | -0.399 – 31.791 |
| 10 | 19 | 12.879 | -0.950 – 30.109 |
| 11 | 20 | 5.521 | -7.274 – 20.403 |
| 12 | 21 | 1.699 | -10.563 – 15.406 |
| 13 | 22 | -0.969 | -13.173 – 12.538 |
| 14 | 23 | -0.914 | -13.471 – 12.566 |
| 15 | 24 | -0.073 | -12.771 – 13.442 |
| 16 | 25 | -0.361 | -13.076 – 13.103 |
| 17 | 26 | -0.700 | -13.331 – 12.885 |
| 18 | 27 | 3.550 | -9.345 – 17.950 |
| 19 | 28 | 3.692 | -8.964 – 17.441 |
| 20 | 29 | 8.950 | -4.332 – 23.713 |

Continued on next page

| Start year | End year | Median (%) | 95% CrI |
|---|---|---|---|
| 21 | 30 | 10.264 | -3.097 – 25.089 |
| 22 | 31 | 11.016 | -2.362 – 26.324 |
| 23 | 32 | 6.123 | -6.937 – 21.394 |
| 24 | 33 | 2.059 | -11.090 – 17.392 |
| 25 | 34 | -0.867 | -13.634 – 13.841 |
| 26 | 35 | -0.233 | -12.432 – 13.856 |
| 27 | 36 | -0.385 | -11.852 – 13.004 |
| 28 | 37 | 1.513 | -9.995 – 14.991 |
| 29 | 38 | 2.680 | -8.840 – 16.459 |
| 30 | 39 | 3.729 | -8.059 – 18.018 |
| 31 | 40 | 7.529 | -4.724 – 22.909 |
| 32 | 41 | 9.958 | -3.545 – 26.399 |
| 33 | 42 | 5.234 | -7.122 – 20.000 |
| 34 | 43 | -1.251 | -12.881 – 11.676 |
| 35 | 44 | -6.900 | -18.576 – 5.539 |
| | | Black Gouse | |
| 1 | 10 | -19.247 | -30.081 – -7.181 |
| 2 | 11 | -16.938 | -27.765 – -5.147 |
| 3 | 12 | -14.080 | -25.001 – -1.847 |
| 4 | 13 | -10.052 | -21.510 – 2.741 |
| 5 | 14 | -8.450 | -20.443 – 4.879 |
| 6 | 15 | -4.828 | -17.147 – 9.375 |
| 7 | 16 | -0.282 | -13.291 – 14.886 |
| 8 | 17 | -1.392 | -14.054 – 13.388 |
| 9 | 18 | -0.474 | -13.335 – 14.474 |
| 10 | 19 | -1.672 | -14.420 – 13.128 |
| 11 | 20 | -5.052 | -17.787 – 9.324 |
| 12 | 21 | -9.164 | -21.660 – 4.387 |
| 13 | 22 | -10.634 | -23.144 – 2.765 |
| 14 | 23 | -9.532 | -22.295 – 4.265 |
| 15 | 24 | -7.977 | -20.296 – 5.230 |
| 16 | 25 | -5.080 | -17.454 – 8.882 |
| 17 | 26 | 1.670 | -11.616 – 17.389 |
| 18 | 27 | 2.558 | -10.826 – 18.706 |
| 19 | 28 | 5.630 | -8.534 – 23.134 |
| 20 | 29 | 5.391 | -8.766 – 23.006 |
| 21 | 30 | 0.977 | -12.427 – 17.281 |
| 22 | 31 | -4.178 | -17.469 – 10.803 |
| 23 | 32 | -7.271 | -20.411 – 7.618 |
| 24 | 33 | -6.254 | -19.677 – 8.825 |
| 25 | 34 | -2.283 | -16.077 – 13.255 |
| 26 | 35 | 3.373 | -10.494 – 19.557 |
| 27 | 36 | 9.994 | -4.395 – 26.987 |
| 28 | 37 | 9.911 | -4.357 – 27.448 |
| 29 | 38 | 13.548 | -1.471 – 32.086 |
| 30 | 39 | 17.749 | 2.038 – 37.884 |
| 31 | 40 | 19.506 | 3.378 – 39.794 |
| 32 | 41 | 17.244 | 1.124 – 36.855 |
| 33 | 42 | 12.492 | -1.935 – 29.739 |
| 34 | 43 | 8.813 | -4.539 – 24.795 |
| 35 | 44 | 5.270 | -7.819 – 20.273 |

### Time series and graphs showing covariates and forest structure

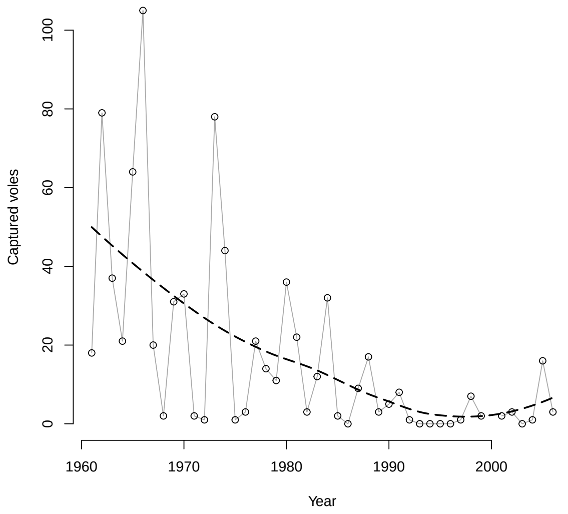


Figure 3: The spring captures of voles in the study area between 1960 and 2024. The stippled line shows the loess function result.

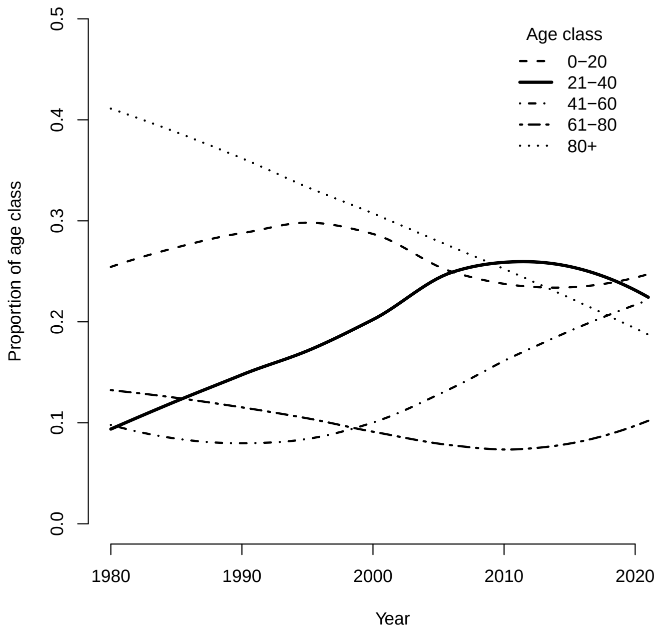


Figure 4: The proportion of forest age classes in the region based on the national forestry inventory[1]. The lines are the results of the loess-function.

[1] *The Swedish National Forest Inventory* (2025). SLU.SE. URL: `https://www.slu.se/en/Collaborative-Centres-and-Projects/the-swedish-national-forest-inventory/about-the-nfi/` (visited on 01/14/2025)

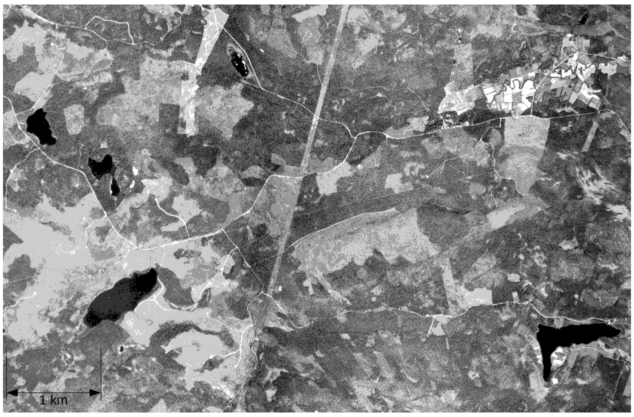


Figure 5: Areal photo of the central parts of the study area in 1975. Light areas are clear-cuts or young plantations.

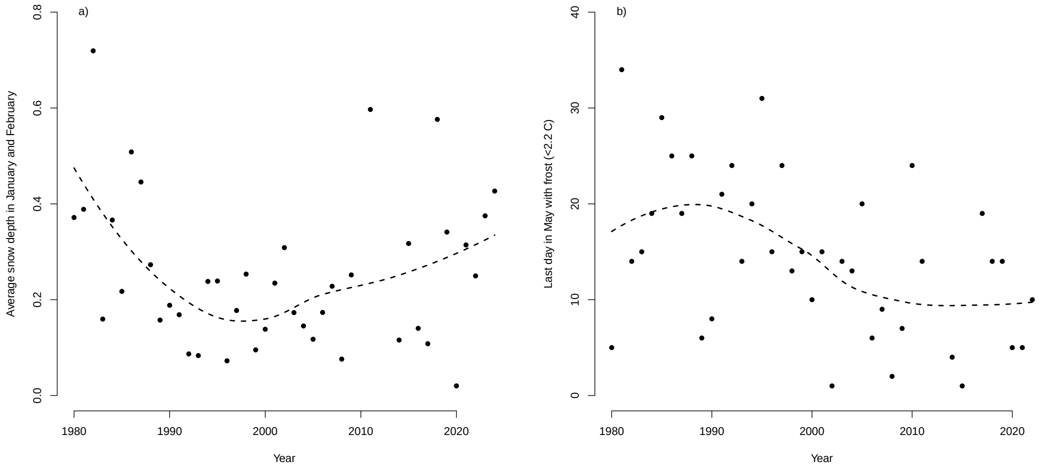


Figure 6: Climate data used in the analysis; a) average snow depth in January and February, b) last day in May with frost (<2.2 C).The stippled lines shows the result of the loess-function.
